# Social interactions are impacted by food availability, food type, and group size

**DOI:** 10.1101/2024.03.07.584014

**Authors:** Xiaohui Guo, Matthew J. Hasenjager, Nina H. Fefferman, Noa Pinter-Wollman

## Abstract

Social interactions are important for how societies function, conferring robustness and resilience to environmental changes. The structure of social interactions can shape the dynamics of information and goods transmission. In addition, the availability and type of resources that are transferred might impact the structure of interaction networks. For example, storable resources might reduce the required speed of distribution and altering interaction structure can facilitate such change. Here we use ants as a model system to examine how social interactions are impacted by group size, food availability, and food type. We compare global- and individual-level network measures across experiments in which groups of different sizes received limited or unlimited food that is either favorable and cannot be stored (carbohydrates), or unfavorable but with a potential of being stored (protein). We found that as group size increased, individuals interacted with more social partners and interaction networks became more compartmentalized. Furthermore, group compartmentalization increased when food was limited and when transferring storable goods. Our findings highlight how biological systems can adjust their interaction networks in ways that relate to their function. The study of such biological flexibility can inspire novel and important solutions to the design of robust and resilient supply chains.

## Introduction

Social interactions facilitate the flow of information, resources, and disease throughout societies. The network structures that emerge from these interactions impact how rapidly and how evenly resources are distributed throughout a population. Both global- and individual-level network features influence these dynamics. For example, networks that are highly modular and subdivided facilitate different flows compared to networks that are uniformly connected (Sah et al., 2017; Salathé and Jones, 2010; Silk and Fefferman, 2021). Similarly, individual connectivity can impact the dynamics of flow on a network, for example highly connected individuals can have a disproportionate impact on disease spread (Lloyd-Smith et al., 2005).

While much theoretical (Eames and Keeling, 2002) and empirical (Stroeymeyt et al., 2014) work has been devoted to uncovering the ways in which network structure influences transmission dynamics, especially in the context of disease transmission (Fefferman and Ng, 2007; Firth et al., 2020; Gates and Woolhouse, 2015; Godfrey, 2013; Jones et al., 2018; Pautasso and Jeger, 2008), less attention has been devoted to the way in which the nature of what is being transmitted might impact network structure. For example, foraging on resources with high spatiotemporal variability may promote food-sharing networks that are structured in a way that mitigates collective resource shortfalls (Jones and Ready, 2022). Likewise, recent theoretical work shows that disease transmission might have shaped the evolution of social interactions (Udiani and Fefferman, 2020) and a recent meta-analysis suggests that the type of disease and its mode of transmission (e.g., air-borne, fluid exchange, etc.) impacts social network structure in animals (Collier et al., 2022). The way in which animals interact is especially important when groups have shared goals, such as social insects, in which sterile workers cooperate to produce reproductives that will found new related colonies. In such cooperative groups, the way in which resources, such as food, are shared, highly depends on how individuals interact with one another (Gordon, 2010). Thus, it is important to determine if the type of resources and their availability impact the structure of social interactions to shape resource sharing in cooperative groups. Here we investigate how the structure of interaction networks that facilitate food distribution respond to the type and availability of resources, in the absence of central control.

Resource availability impacts how individuals interact. For example, when carbohydrates are limited, animals increase their foraging for sugars (Hendriksma et al., 2019; Kay, 2004) and similarly, they increase their intake of proteins and other macronutrients when those are limited (Kohl et al., 2015; Mayntz et al., 2005). Limitations on resources may alter how they are shared among group members and therefore such limitations can impact the way in which individuals interact. Abundant food resources reduce the need for food-sharing if each individual is able to supply itself adequately, resulting in fewer interactions overall. Similarly, if food supply is limited, we might still expect few interactions if sharing a scarce resource means that each group member will not receive enough to survive. Distributing scarce resources within smaller subgroups may ensure that at least some individuals receive sufficient resources to survive until resources availability increases, at the potential cost of losing some group members that are in subgroups that do not receive enough resources. Thus, if food availability influences interaction structure, we might expect more subdivided networks when food is scarce and interaction networks with a single or very few clusters when food is abundant.

Furthermore, the type of resources that are distributed can influence the way in which individuals interact. Some resources can be stored and their dissemination does not rely on rapid sharing, while perishable resources that cannot be stored may need to be distributed rapidly. Animals utilize different types of nutrients to serve different physiological needs, which can change over time (Simpson and Raubenheimer, 2012). For example, when offspring are being produced, ants require and forage for more proteins than when they do not have offspring to feed (Cassill and Tschinkel, 1995). Thus, it is possible that the type of resource which is being shared in a group might determine how it is shared, altering network structure. For example, the presence of perishable goods may result in more interactions to facilitate rapid resource dissemination, compared to when storable resources are being distributed.

Finally, group size is an important factor that shapes patterns of social interaction. The more individuals in a group, the greater the potential to interact (Quque et al., 2021). However, increased interactions can be costly—e.g., individuals may be at greater risk of exposure to pathogens and injury due to aggression. One way to mitigate such costs may be to subdivide large groups into clusters in which interactions occur more intensely compared to across clusters (Silk and Fefferman, 2021). Thus, as a group becomes larger, global changes to group structure, or to the way in which a group is organized, may alter how individuals interact (Miller et al., 2022). Groups of different sizes may have different interaction patterns that maintain certain network features (O’Donnell and Bulova, 2007; Pacala et al., 1996). For example, if the number of interactions increases with group size, network density (the number of observed connections over the number of possible connections) may remain constant across group sizes if network density is important for the robustness of resource flow. Alternatively, if the number of interactions does not scale with group size, network density may decrease with group size, potentially slowing down the transfer of goods in larger groups. Here we ask how group size impacts the structure of a network that is used to transfer resources.

Ants are an ideal model system for examining the dynamics of social interactions because of the important function that interactions have for the fitness of the colony. The way in which ants interact with one another allows them to regulate their collective foraging (Gordon, 2010; Greene and Gordon, 2006; Greene et al., 2013; Pinter-Wollman et al., 2013) for example in response to colony nutritional needs (Csata et al., 2020; Dussutour and Simpson, 2008; Dussutour and Simpson, 2009). Certain individuals (i.e., foragers) leave the nest to collect food and bring it back to the nest (Gordon, 1989; Gordon, 1996). Once at the nest, food is distributed and stored, and foragers decide whether or not to continue foraging based on certain types of interactions with nestmates (Miller and Pinter-Wollman, 2023), the forager’s own food load (Greenwald et al., 2018; Howard and Tschinkel, 1980; Wallis, 1964), how deep a forager moves into the nest (Baltiansky et al., 2023), and the presence of larvae in the nest (Ulrich et al., 2016).

The sharing of liquid food, which many ant species forage for and consume, is carried out through trophallaxis, which is a mouth-to-mouth interaction in which liquid food is transmitted from one individual to another. Resource availability impacts the speed of food dissemination through the colony: the longer a colony is starved, the quicker newly discovered food is distributed throughout the colony (Howard and Tschinkel, 1981). This change in food distribution speed might be a result of changes to the interaction network that facilitates food sharing (Sendova-Franks et al., 2010). Furthermore, ant colonies require different types of nutrients, with workers primarily consuming sugars, and proteins being consumed by queens and larvae (Markin, 1970). Proteins have a negative impact on worker longevity (Dussutour and Simpson, 2012) and ant foraging decisions are impacted by the type of food they require (Barbee and Pinter-Wollman, 2022; Portha et al., 2002). Because sugars are consumed by workers, they can be viewed as a perishable resource that is not stored, while proteins can be stored in the brood. Sugars are distributed faster among workers than proteins (Howard and Tschinkel, 1981), potentially because ants have more interactions when fed with sugars than when fed with protein. Finally, most food transmission inside the nest occurs among workers, rather than to the queen or larvae, (Wilson and Eisner, 1957) and the number of trophallaxis interactions can increase with group size (Quque et al., 2021). Therefore, group size can impact the way in which food is distributed by altering interaction patterns. Given the ways in which ants respond to food availability, the array of nutritional needs within the colony, and the importance of group size for interactions, ants are an excellent system for examining how the nature and utility of different resources may shape the ways in which resources are distributed.

By studying the interactions among carpenter ants (*Camponotus fragilis*) under different conditions we ask if social interactions are impacted by group size, food availability, and food type. We quantify interactions using both global- and individual-level network measures and predict that larger groups will have more interactions to facilitate the transfer of resources. Furthermore, when food supply is unlimited, we expect fewer interactions than when supply is limited if distribution is less important for accessing resources. However, when food supply is limited, we might also expect a decrease in the number of interactions if it is suboptimal to share the scarce resource throughout the entire group (i.e., if each group member will not receive sufficient resources). However, clustering may increase when supply is low relative to when it is unlimited to ensure that at least some group members receive resources. Finally, we expect that when preferable food sources are provided (carbohydrates for workers), there will be more interactions and less subdivision of the network (i.e., fewer clusters), compared to when a less preferred resource is provided (protein), to expedite the flow of the preferred resource.

## Methods

### Ant maintenance, tagging, and preparation for experiments

We obtained worker ants of the species *Camponotus fragilis* from a colony maintained by an ant supplier (John Truong) on September 7, 2021. After transfer to the lab at UCLA, ants were housed in a rectangular plexiglass container (10.16 cm x 5.08 cm x 2.54 cm) and fed twice a week with liquid protein-rich food and carbohydrate-rich food (see recipes in the Supplemental materials). We tagged ants with BEEtags (Crall et al., 2015) that were printed on paper and laminated using transparent Scotch® tape. We used Loctite epoxy adhesive to affix the tags to the ants’ thorax (Figure 1B). Before each experimental trial, tagged ants were selected haphazardly and placed together as a group in a petri dish (90×15mm) for 5-6 days without food. Group sizes ranged from 14-30 individuals. All experiments were conducted between October 26, 2021 and June 24, 2022.

**Figure 1:**
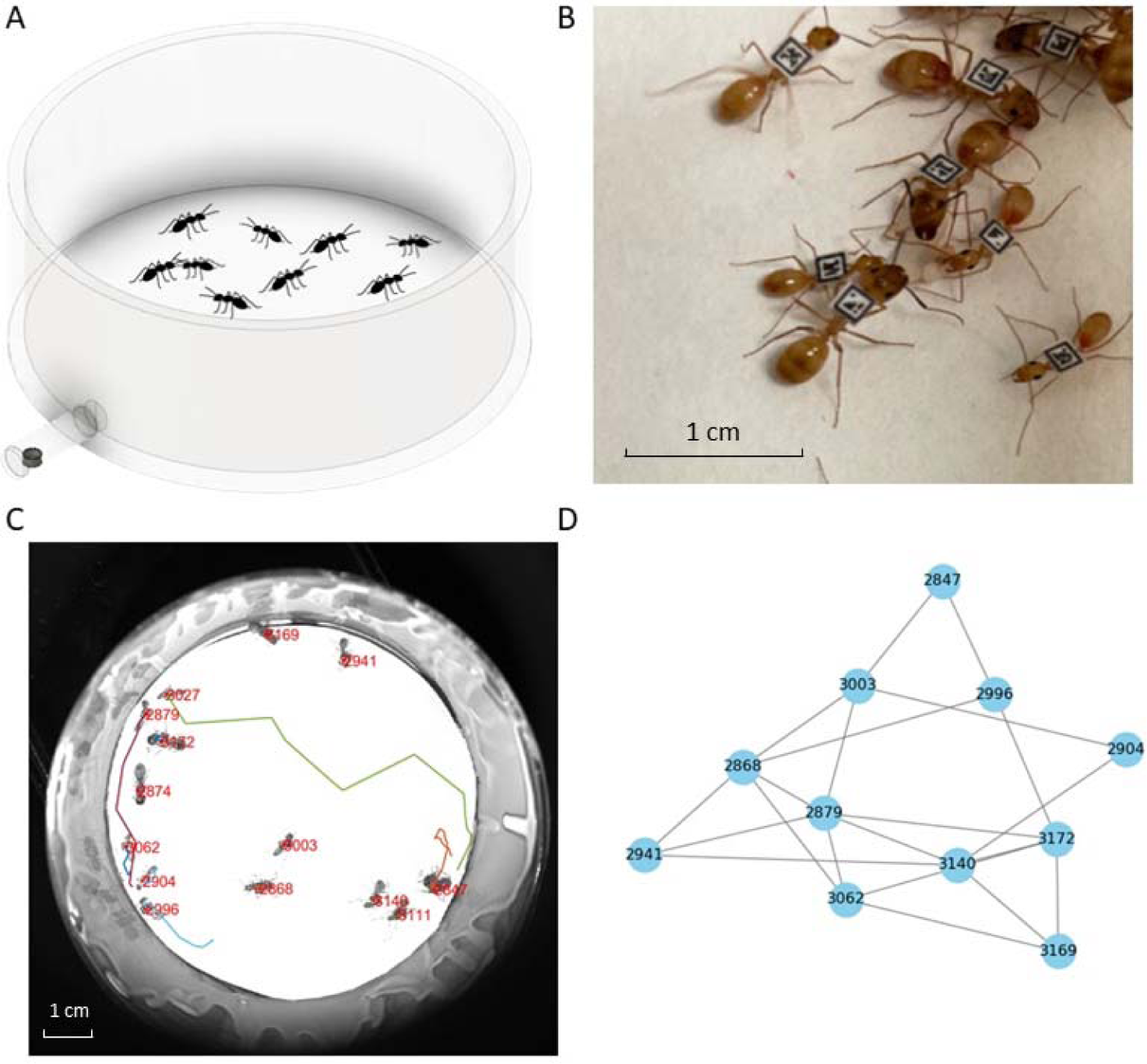
Experimental setup. A. Illustration of the petri dish in which ants were tracked during the experiments, with the tube that led to the food source at the bottom left (ants not to scale). B. Image of ant workers (*C. fragilis*) with individually attached BEEtags. C. An image from one of the experiments overlaid with ants’ trajectories and their individual IDs - as identified from their BEEtags. D. A social network inferred from imaging data; nodes are individual ants that are connected with edges if their heads came close enough to one another to allow for trophallaxis.

### Experimental treatments

Each group of ants was assigned haphazardly to one of four treatments in which we provided them with either an unlimited or a limited supply of food that was either rich in carbohydrates or in protein. During the experiments, we supplied ants with food outside the petri dish in which they were housed and tracked (see tracking details in the following section). Liquid food (0.3 ml) was placed in a cap of a microcentrifuge tube which was connected to the petri dish with a plastic tube (inner-diameter = 5mm; length = 2 cm;) that allowed access to the food by only one ant at a time (Figure 1A).

#### Unlimited food supply

When food supply was unlimited, we provided the food, as detailed above, for the duration of the experiment. We provided enough food so that it did not run out and did not require replenishing during the experiment.

#### Limited food supply

When food supply was limited, we provided the ants with food as detailed above until 10% of the unique ants in the group visited the food source, fed, and returned to the tracked petri dish. Once 10% of the ants fed (this took less than 5 min), we closed the tube connecting the tracked petri dish to the food using a cotton ball. The time it took 10% of the ants to visit the food was so short that no ant visited the food more than once.

#### Carbohydrate-rich food

We made carbohydrate-rich food by mixing 0.22g sugar and 2 ml deionized water.

#### Protein-rich food

We made protein-rich food by mixing 0.564g pasteurized egg powder (Modernist Pantry), 0.188g sugar and 3.75ml deionized water.

Both recipes are modified from (Baltiansky et al., 2021). Because workers died during the course of the study, especially when fed with protein, as seen in other studies of ant diet (Dussutour and Simpson, 2012), our experimental design was not completely balanced. For sample sizes in each treatment, see Table 1.

**Table 1:** Sample sizes (number of groups tested) in each treatment combination.

| Food type: | Carbohydrates | Protein |
| --- | --- | --- |
| Food availability: |  |  |
| *Unlimited* | 13 | 8 |
| *Limited* | 13 | 12 |

### Filming, image analysis, and inferring interactions

Groups were filmed for 60 min using a camera (FLIR Blackfly) and mounted LED lights for illumination (Thorlabs, MCWHL6) and images were captured using the Micromanager software (Edelstein et al., 2014). Because of the time it took to store images while filming, the frame rate was not constant (it slowed down over time) and ranged from 0.31 to 2.92 images/sec. Because we did not study the duration of interactions, only whether or not an interaction occurred, this slight variation in frame rate was not accounted for in our analysis. The position of each ant was extracted from the individually attached BEEtags using a slightly modified BEEtag tracking code in Matlab (Crall et al., 2015) (Figure 1C), code available on Github (https://github.com/MJHasenjager/Identifying-Trophallaxis-Networks). We used the directionality of the BEEtags to determine the position of each ant’s head and considered only head-to-head interactions in our analysis. Considering only head-to-head interactions allowed us to restrict our analysis to interactions that might result in trophallaxis (i.e., exchange of liquid food) and exclude other interactions, such as head-to-abdomen, abdomen-to-abdomen, etc. To identify interactions, we first determined manually for each trial the distance between ants’ heads while they were interacting in 50 randomly selected frames. We then used the maximum manually measured distance as the threshold distance to automatically infer interactions throughout the trial. If the heads of two ants were equal to or less than this threshold distance in any frame, they were considered interacting. Because a different threshold distance was used in each trial, this distance ranged from 46 to 102 pixels (1pixel = 0.04mm), code available on Github (https://github.com/MJHasenjager/Identifying-Trophallaxis-Networks). Due to variation in frame rates across and within trials, we did not determine the strength (duration) of each interaction, but simply noted if individuals interacted or not to form an unweighted and undirected interaction network. Note that not all ants interacted and therefore interaction networks might include fewer individuals than the number of individuals in a group (Figure S1).

### Social network analysis

We used network analysis to quantify the social behavior of the ants and to determine how their social behavior changed in response to the experimental manipulations. Each ant was a node in the network, and an interaction (edge) connected two ants when their heads were within a specified distance threshold that allowed for trophallaxis, as detailed above (Figure 1D). To quantify the social behavior of the ants we used two network measures that quantify global network structure (network density and number of clusters) and two individual-based centrality measures (degree and betweenness). Network measures were calculated using the R package ‘igraph’ (Csardi and Nepusz, 2006).

#### Network density

Number of observed links among ants divided by all possible connections. This measure provides information about the overall connectivity of the network while scaling for network (group) size.

#### Number of clusters

Number of clusters that each network can be delineated into. We applied the *‘walktrap’* clustering algorithm (using the cluster_walktrap() function in the R package ‘igraph’ (Csardi and Nepusz, 2006)) to all networks and recorded the number of clusters that were identified by this algorithm. We selected the *‘walktrap’* algorithm, among many available network clustering algorithms, because on visual inspection of the clusters that it identified, it provided the most biologically plausible clusters, with breaks between clusters where groups were least connected.

#### Degree

Number of unique individuals that an ant interacted with. Provides information on how many interaction partners each individual had. This measure is not scaled for group size and so there are often more opportunities to have greater degree values in larger groups.

#### Betweenness

Number of shortest paths that connect pairs of individuals and that pass through the focal ant. This measure provides information on how well each ant acts as a bridge for other ants’ interactions. Individuals with high betweenness connect many ants which might not be well-connected themselves.

### Statistical analysis

To determine the impact of group size, food availability, and food type of social interactions, we ran linear mixed models (LMM) or generalized linear mixed models (GLMM). In each model one of the network measures (density, number of clusters, degree, or betweenness) was the dependent variable. The explanatory variables included group size (a continuous numeric value), food availability (limited or unlimited), and food type (carbohydrates or protein). For group-level network measures, each data point was for an entire group and we included ‘group ID’ as a random effect because some groups were used more than once in our experiments. For individual-based measures, each data point was for an individual ant, and we included both ‘group ID’ and ‘individual ID’ as random effects in the model to account for variation among groups and among individuals that were tested multiple times in different treatments.

We used a model selection approach to determine whether or not to include interactions among effects in our final statistical model. We ran each model with either no interactions among group size, food type, and food availability; with the three-way interaction term among the three variables; and three additional models with just one interaction each between a different pair of variables each time, totaling 5 statistical models. We then compared the models using AIC and selected the best fit model, i.e., the one with the lowest AIC score. The best fit models for the two global-level measures, density and number of clusters, included no interaction terms among the explanatory effects. The best fit models for the two individual-level network measures included all possible interaction terms among the three fixed explanatory effects (see supplementary materials for AIC values of all the models we tested). All the best fitting models met the required statistical assumptions – examined using the check_model() function in the ‘performance’ package (Lüdecke et al., 2021).

For density, number of clusters, and degree, we ran an LMM, implemented using the lmer() function in the ‘lme4’ package. For betweenness, we used a GLMM with a gamma log link function, implemented using the glmer() function in the ‘lme4’ package (Bates et al., 2015). We report the analysis of deviance of the models, obtained using the Anova() function in the ‘car’ R package (Fox and Weisberg, 2019). We report the percent variance explained by the random effects as the conditional R^2^ minus the marginal R^2^. For the two final models of the individual-level network measures, we conducted post hoc Tukey tests using the emmeans() function in the ‘emmeans’ R package (Lenth, 2022).

Image analysis was conducted in Matlab (Mathworks Inc., Natick, MA, U.S.A.) and network and statistical analyses were conducted in R (R Core, 2014).

### Data accessibility

All data and code can be found on Github (https://github.com/MJHasenjager/Identifying-Trophallaxis-Networks).

### Ethical Note

This work was conducted in accordance with the local animal welfare laws, guidelines and policy for the use of animals in research. Ants are invertebrates and do not require special institutional permissions for experimentation. We handled ants with extreme care. We used soft tweezers when handling the ants to minimize harm. Experiments involved video recording of ants’ behavior, with no invasive methods. After the experiments we kept the ants in the lab and provided them with food *ad lib* until they died naturally

## Results

Group size, food type, and food availability differed in their impact on the four measures of social behavior. Of the group-level measures, only the number of clusters was impacted by our experimental treatments, but both individual-level centrality measures (degree and betweenness) were highly impacted by the different experimental manipulations.

Network density was not impacted by group size, food type, or food availability. None of the effects in the model were statistically significant (Table 2). The random effect ‘group ID’ explained 17% of the variance in the model.

**Table 2:** Analysis of deviance (Type II Wald *X*^2^ tests) for the LMM examining the effect of group size, food type, and food availability on network density.

| Effect | $X^2$ | DF | p-value |
| --- | --- | --- | --- |
| Food type | 1.886 | 1 | 0.169 |
| Food availability | 1.882 | 1 | 0.170 |
| Group size | 0.013 | 1 | 0.911 |

The number of clusters in a network was significantly impacted by group size, food type, and food availability (Table 3). Larger groups had significantly more clusters (Figure 2A, Table 3). Furthermore, when provided with protein-rich food there were significantly more clusters than when provided with carbohydrates-rich food (Figure 2B, Table 3). Finally, as predicted, when food was unlimited there were significantly fewer clusters than when food was limited (Figure 2C, Table 3). The random effect ‘group ID’ explained 12.1% of the variance in the model.

**Figure 2:**
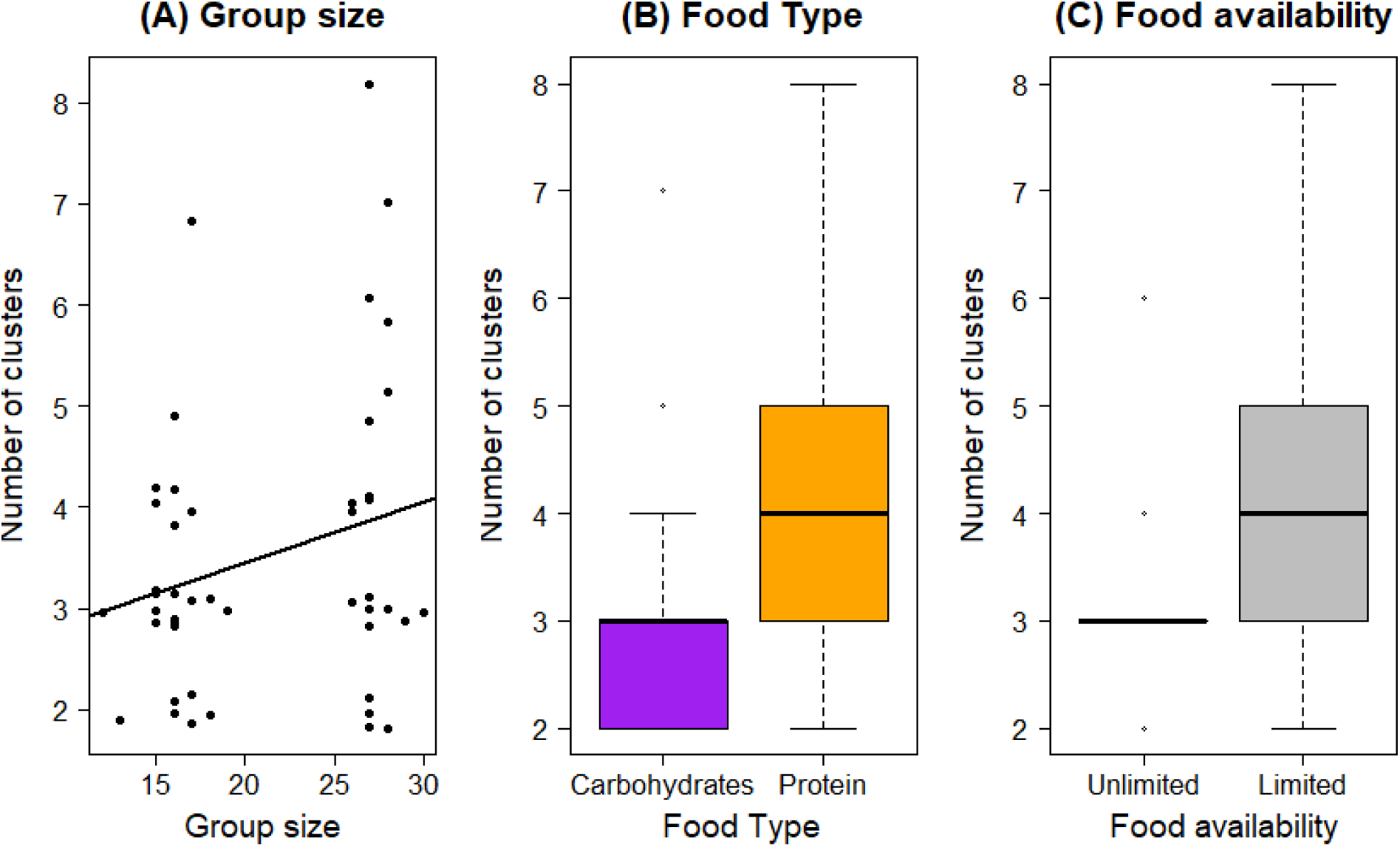
Number of clusters in the interaction network was impacted by group size (A), food type (B), and food availability (C). In these and all other boxplots, the black horizontal line is the median, boxes extend to 25 and 75 percentiles, whiskers extend to 1.5 times the interquartile range, and points are outliers. In (A) the points are slightly jittered along the y-axis to improve visibility and the line shows the model fit.

**Table 3:** Analysis of deviance (type II Wald ^2^ tests) for the LMM examining the effect of group size, food type, and food availability on number of clusters in a group. Bold p-values indicate statistically significant results.

| Effect | <sup>2</sup> | DF | p-value |
| --- | --- | --- | --- |
| Food type | 7.725 | 1 | **0.005** |
| Food availability | 5.935 | 1 | **0.015** |
| Group size | 4.136 | 1 | **0.042** |

The number of unique individuals that each ant interacted with (i.e., degree) was significantly impacted by group size, food type, food availability, and all the interactions among these variables (Figure 3, Table 4). As expected, ants in larger groups interacted with more individuals. Interestingly, ants in groups that were fed with carbohydrate-rich food interacted with significantly more unique individuals than ants in groups fed with protein-rich food (Figure 3, Table 4). When examining the interaction between the effect of group size on degree and food type, we found that the positive association between degree and group size was enhanced when ants were fed with carbohydrate-rich food compared to when fed with protein-rich food (post hoc Tukey test comparing slopes by food type: df = 82.4, t = 2.02, p = 0.0466). Similarly, the positive relationship between degree and group size was stronger when the food supply was unlimited compared to when it was limited (post hoc Tukey test comparing slopes by food availability: df = 710, t = 5.911, p < 0.0001). When examining the interaction between all three fixed effects with post hoc tests, we found that when groups were fed with carbohydrate-rich food and food was unlimited, degree increased to a greater extent with group size compared to when ants were fed with a limited supply of carbohydrate-rich food (post hoc Tukey test comparing slopes: df = 553, t = 7.254, p < 0.0001) or when ants were fed with limited supply of protein-rich food (post hoc Tukey test comparing slopes: df = 75.7, t = 4.326, p < 0.0003).

**Figure 3:**
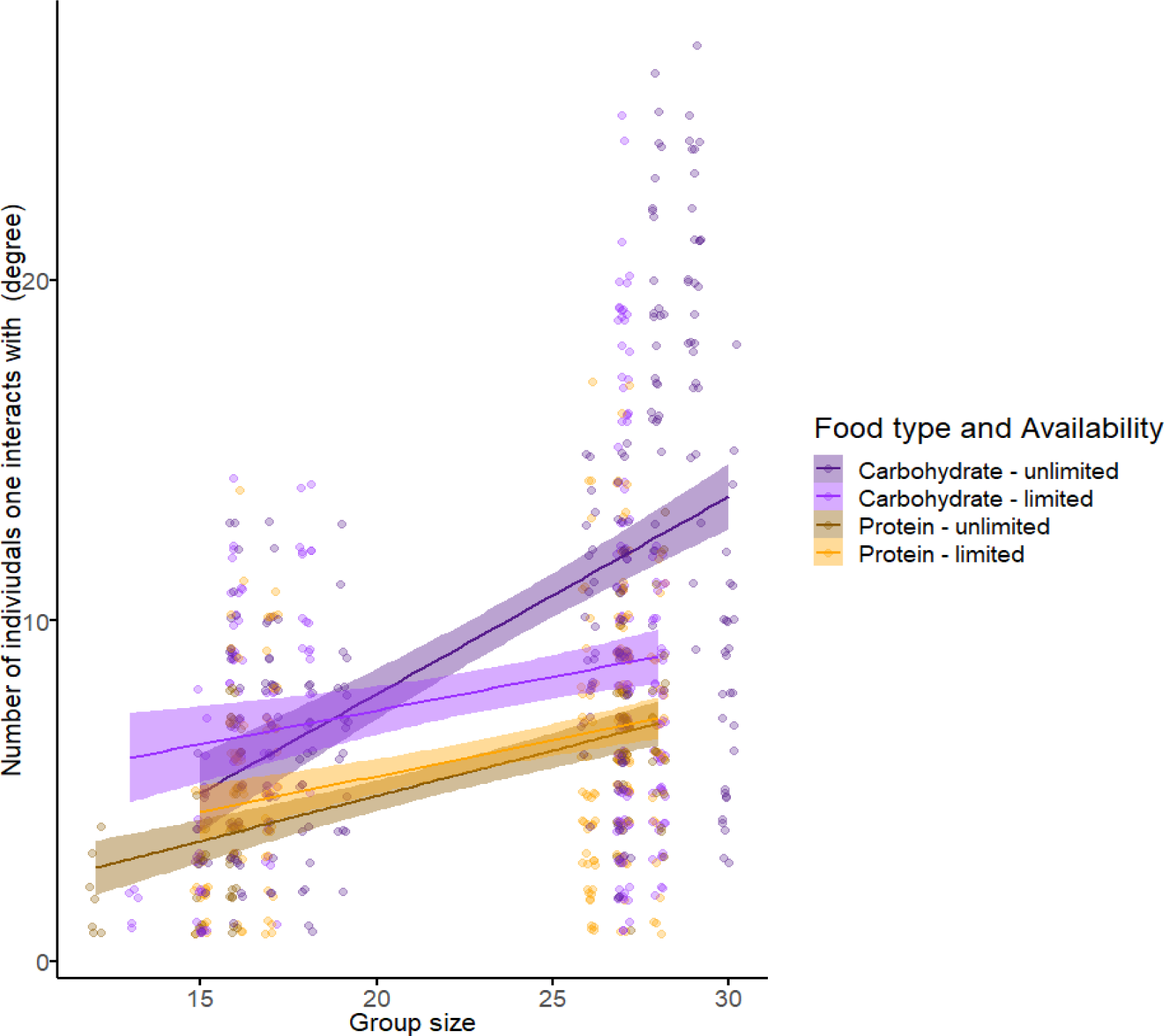
Number of unique individuals that an ant interacted with (degree) was significantly impacted by group size, with different relationships by food type and food availability. Each point represents an individual ant. Points in purple are from experiments in which the food type was carbohydrate-rich and orange points are from experiments in which the food type was protein-rich. Darker and lighter colors respectively denote experiments in which food supply was unlimited and limited. Lines show the model fit with shaded areas as 95% confidence intervals. Points are slightly jittered along the x and y axes to improve visibility.

**Table 4:** Analysis of deviance (type II Wald *X*^2^ tests) for the LMM examining the effect of group size, food type, food availability, and the interactions among them on the number of unique individuals each ant interacted with (degree). Bold p-values indicate statistically significant results.

| Effect | $X^2$ | DF | p-value |
| --- | --- | --- | --- |
| Food type | 33.064 | 1 | < **0.0001** |
| Food availability | 38.107 | 1 | < **0.0001** |
| Group size | 37.580 | 1 | < **0.0001** |
| Food type x Food availability | 6.575 | 1 | **0.01** |
| Food type x Group size | 5.425 | 1 | **0.02** |
| Group size x Food availability | 25.720 | 1 | < **0.0001** |
| Food type x Group size x Food availability | 109.898 | 1 | < **0.0001** |

However, the increase in degree with group size when fed with unlimited supply of carbohydrate-rich food was not significantly different compared to when ants were fed with an unlimited supply of protein-rich food (post hoc Tukey test comparing slopes: df = 105, t = 2.237, p = 0.120). The random effects ‘group ID’ and ‘individual ID’ explained 41.7% of the variance in the model.

The number of shortest paths between pairs of ants that pass through a focal ant (i.e., betweenness) was significantly impacted by group size, food availability, and the interaction between them, but not by food type (Figures 4 & 5, Table 5). The only effect of food type on betweenness was through its interaction with food availability. When food was limited, betweenness was greater than when food was unlimited (Table 5, Figure 4A). This difference was driven by trials in which ants were fed with protein-rich food, but not by trials when ants were fed with carbohydrate-rich food. When ants were fed predominantly carbohydrates, there was no statistically significant difference in betweenness when comparing trials in which food was limited compared to those with unlimited food (Figure 4B, post hoc Tukey test comparing food availability within carbohydrate food type: Z ratio = −0.633, p-value = 0.9215). However, when ants were fed with protein-rich food, there was a statistically significant difference in betweenness when comparing trials in which food was limited compared to when food was unlimited (Figure 4C, post hoc Tukey test comparing food availability within protein food type: Z ratio = −3.240, p-value = 0.0066). Finally, the positive relationship between betweenness and group size was impacted by food availability, with the slope being significantly greater when food was limited compared to when it was unlimited (Figure 5, post hoc Tukey test comparing slopes by food availability: Z ratio = −2.978, p-value = 0.0029).

**Figure 4:**
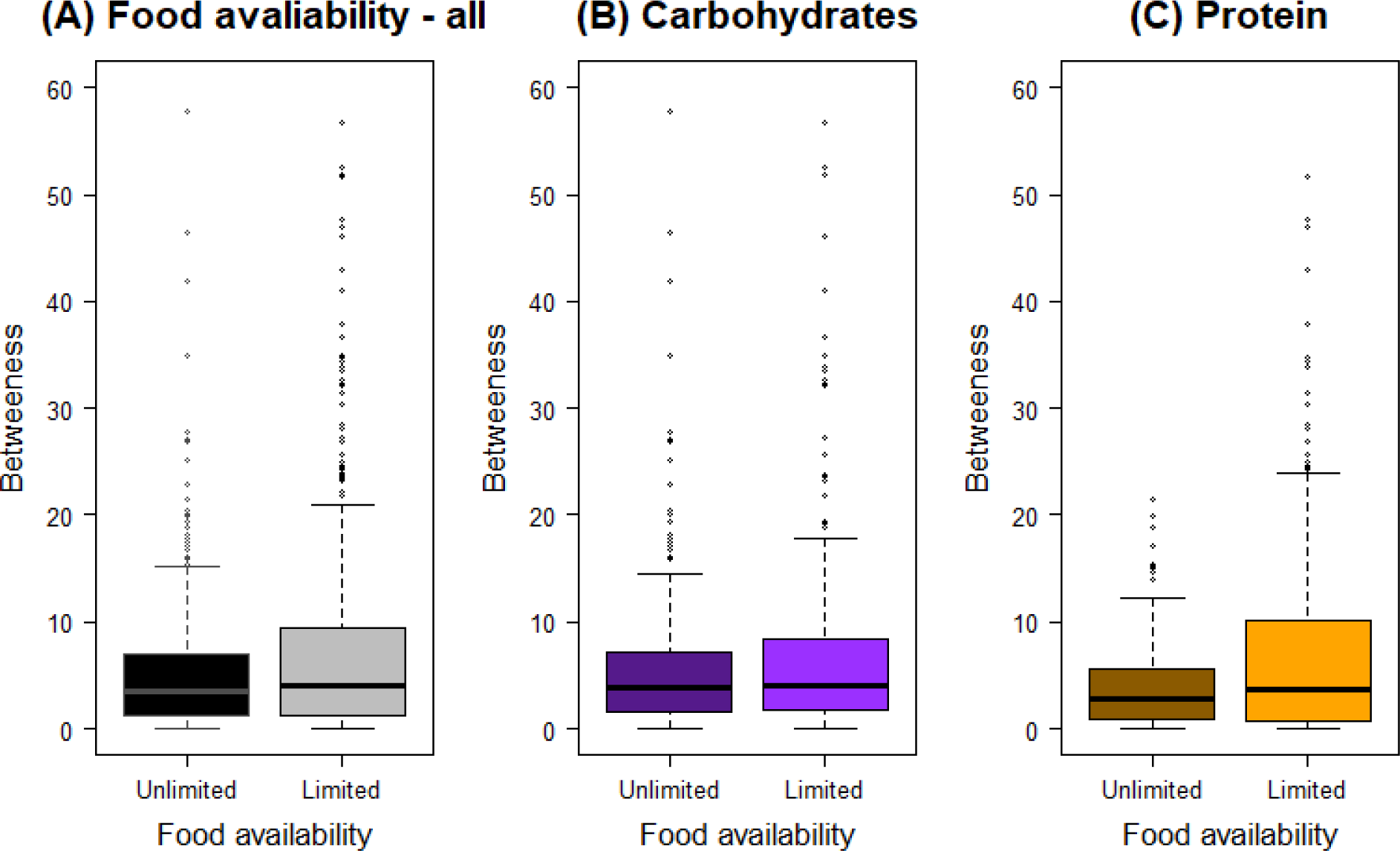
Betweenness was impacted by food availability differently when ants were fed with proteins or with carbohydrates. (A) Betweenness was greater when food supply was limited (gray) than when it was unlimited (black). (B) When ants were provided with a preferred food source (carbohydrates), there was no effect of food supply on betweenness. However, when ants were provided with a less preferred food (protein) (C), betweenness increased when food supply was limited.

**Figure 5:**
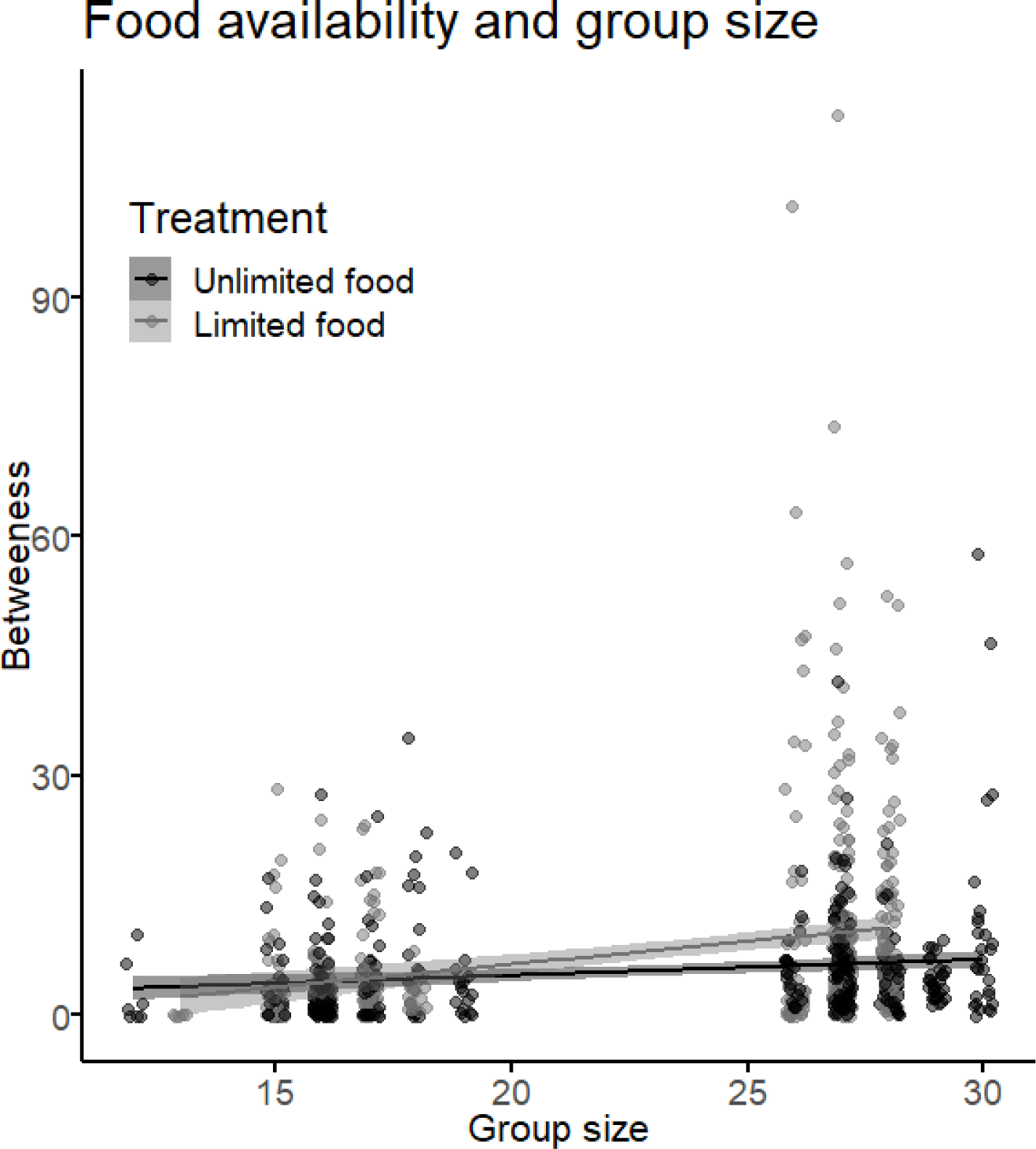
When food supply was limited (gray), betweenness increased to a greater extent with group size than when food was unlimited (black). Points are slightly jittered along the x axis to improve visibility and lines are the fit of the statistical model with shaded areas as the 95% confidence intervals.

**Table 5:** Analysis of deviance (type II Wald *X*^2^ tests) for the statistical model examining the effect of group size, food type, food availability, and the interactions among them on the number of shortest paths that pass through a focal ant (betweenness). Bold p-values indicate statistically significant results.

| Effect | $X^2$ | DF | p-value |
| --- | --- | --- | --- |
| Food type | 0.035 | 1 | 0.852 |
| Food availability | 6.728 | 1 | **0.009** |
| Group size | 40.955 | 1 | **< 0.0001** |
| Food type x Food availability | 5.556 | 1 | **0.018** |
| Food type x Group size | 0.325 | 1 | 0.568 |
| Group size x Food availability | 7.719 | 1 | **0.005** |
| Food type x Group size x Food availability | 2.116 | 1 | 0.146 |

## Discussion

We found that ant social interactions were impacted by group size, food availability, and food type. As one might expect, the number of unique individuals contacted by each ant and the number of social clusters in a colony both increased with group size. Interestingly, both limiting food supply and providing a less preferred, potentially storable, food source resulted in more clustered interaction networks and in fewer interactions with unique individuals. These observed changes to network structure have the potential to impact the rate of food transfer among ants because the head-to-head interactions that we observed reflect the potential to engage in food-sharing behaviors.

Individuals in larger groups interacted with more unique individuals (high degree, Figure 3) and we detected more clusters in larger groups (Figure 2A). However, network density, i.e. the observed number of links relative to the possible number of links, did not increase with group size. Because network density is scaled to group size, we can infer that the positive relationship between other network measures and group size emerges simply from having more ants in the group and not from qualitative changes in the way that ants interact when they are in larger groups. The petri dishes used to house ants in all experiments were the same size, so ants in larger groups were more crowded. Despite this greater crowding of ants in larger groups, network density did not change with group size, suggesting that ants maintained a relatively constant interaction rate across population densities (i.e., number of ants per unit area) in the range that we tested. A study of the trophallaxis interactions of the black garden ant similarly found that the distribution of food throughout the colony is independent of group size (Quque et al., 2021). One way to decouple group size and network density is by reducing the number of ants that participate in interactions. However, we found that most (but not all) ants in a group participated in the interaction networks that we observed (Figure S1). Only when ants were fed with an unlimited supply of protein-rich food did the proportion of ants that participated in the interaction network slightly decline relative to other treatments (Figure S2). In addition, maintaining interaction network density across group sizes can be achieved by increasing the number of interactions that each individual experiences with group size. Indeed, we found that as group size increases, ants interact with more unique individuals (Figure 3), similar to the positive relationship between number of interactions and group size in black garden ants (Quque et al., 2021). Thus, by adjusting the local behavior of each ant (degree), the global structure of the network (network density) is maintained across group sizes and across population densities.

Food availability impacted both global- and individual-level interaction measures. When food was limited, the ants formed more clusters than when food was not limited (Figure 2C). Similarly, the individual-level centrality measure betweenness increased when food was limited compared to when it was unlimited (Figure 4A), particularly within groups that were fed with protein-rich food (Figure 4B, C). More clustering means that there is greater subdivision within an interaction network, increasing the potential for certain individuals to act as ‘brokers’ among clusters, i.e., having higher betweenness. Similarly, work on *Formica fusca* ants showed that certain individuals in the colony can accumulate food, essentially acting as food storage units that can enhance food distribution across the network (Buffin et al., 2009). While having many clusters may slow down food transfer throughout the entire group, it can also expedite food sharing within clusters, because there are fewer individuals within each cluster (Sah et al., 2017). These dynamics may result in homogeneous food distribution within, but not across, clusters. When resource supply is limited, such a distribution of interactions might be advantageous to the entire group as a bet hedging strategy, helping to ensure that at least part of the group (e.g., all individuals in one or more clusters) may have access to a sufficient amount of resources. Without such social sub-division, a limited resource might be distributed evenly across the entire group, risking a situation in which everyone receives too little food to survive. Indeed, within human populations that rely on subsistence harvesting, it has been argued that a modular organization of the population into local resource-sharing clusters can help to mitigate the impacts of systemic resource shortfalls by limiting local collapses (Jones and Ready, 2022). While trophallaxis interactions facilitate a homogenous distribution of cuticular hydrocarbons which are critical for nestmate recognition (Dahbi et al., 1999), a homogeneous distribution of food to all nestmates might not always be necessary, or advantageous. Our finding that betweenness increased with group size when food was limited but not when it was unlimited (Figure 5) further suggests that subdividing an interaction network may be important for preventing the dilution of a resource across all group members, especially when supply is limited and the population is large.

Food type impacted both global- and individual-level network measures, with parallels to some of the results we saw for food availability. Ants formed more interaction clusters and interacted with fewer other ants when fed with protein-rich food compared to when they were fed with carbohydrate-rich food. Thus, they responded to protein in a similar manner to their response to limited food availability. The greater degree and stronger positive relationship between degree and group size that we observed when feeding ants with carbohydrate-rich food could reflect a higher demand for carbohydrates by workers. Indeed, ant workers consume and use carbohydrates for energy, whereas protein is transferred to the brood and queens (Markin, 1970). However, while brood are the main consumers of protein (Howard and Tschinkel, 1981), when providing ants with protein in liquid form mixed with sugars, similar to the protein-rich food used in our study, such food can remain in workers for 24 hours (Sorensen and Vinson, 1981). Furthermore, the lack of brood in our experimental groups might have resulted in decreased need to distribute protein-rich food throughout the group, however, previous work found that the presence of brood does not impact the way in which ants interact (Quque et al., 2021). Finally, carbohydrates cannot be stored and are akin to perishable goods, whereas protein can be stored in larvae and is thus a potential proxy for non-perishable goods. Differences in storability of goods may influence individual- and system-level behavior of supply networks. Perishable goods may need to be distributed rapidly throughout the population, enabled by a network with few clusters and high node degree. However, when goods can be stored, speed of distribution may be less important, favoring other forms of network organization such as the formation of clusters that specialize in acquiring and sharing certain storable goods, thereby freeing-up other network clusters to perform other activities, which might increase organization efficiency.

Further work is needed to uncover the mechanisms that underlie the impacts we found of group size, food availability, and food type on interaction patterns. For example, changes to the level of activity of workers and their walking patterns when fed different types of food might explain changes in interaction rates (Pinter-Wollman, 2015b). Furthermore, the spatial distribution of ants within the nest might determine who interacts with whom and how frequently (Pinter-Wollman, 2015a; Pinter-Wollman et al., 2013; Pinter-Wollman et al., 2011) and can influence foraging decision and the flow of food among nestmates (Baltiansky et al., 2023; Buffin et al., 2009). It might be interesting to determine who initiates interactions, fed or hungry ants because recent work suggests that fed and hungry ants play different roles in food distribution within ant colonies (Miller and Pinter-Wollman, 2023). In addition, our work did not distinguish between different types of interactions, for example, when ants interact head-to-head, they might be antennating, connecting mandibles without food exchange, or actively exchanging food. Furthermore, ants may have other types of encounters, such as head to body – which may lead to exchange of information through cuticular hydrocarbons, but not to food exchange. While we did not distinguish between different types of interaction, recent work shows that trophallaxis interactions are more important than other interaction types in determining ant foraging decisions (Miller and Pinter-Wollman, 2023). While our work was limited to relatively small groups of ants without brood or queens, the fact that we detected such a large impact of food type and availability on interaction patterns suggests that it is important to consider the type and availability of goods when studying supply chain structure and dynamics. It is possible that some of the patterns we observed here are buffered in large colonies that contain brood. For example, if only a small proportion of the colony participates in interactions to distribute food, the number of individuals engaged in food distribution might be constant and social interactions that facilitate food distribution would not be impacted by the effects of group size that we found here. Furthermore, the presence of brood that can store (and consume) proteins as well as the negative effects that a protein-rich diet might have on the survival of workers (Dussutour and Simpson, 2012), might further impact the way in which workers interact when provided different types of food. Finally, ant species differ in their propensity to transfer liquid food among workers (Wilson and Eisner, 1957) and a comparative study of food transmission dynamics across ant species might reveal a variety of transmission strategies that are adapted to different environments.

To conclude, our work provides insights into how social interactions that can facilitate the exchange of goods change in response to group size and to the availability and type of goods. Interactions increase and become more compartmentalized as groups size increases, as supplies become limited, and when transferring storable goods. However, when perishable goods with high demand are being transferred, we see many interactions that increase rapidly with group size and are not compartmentalized. This adjustment of biological social networks to potentially improve their function may inspire the study and design of robust and resilient supply chains.

## Acknowledgments

We would like to thank John Truong for the ant supply. Sean O’Fallon, Kaija Gham, and Karen Mabry provided helpful suggestions during the preparation of this manuscript.

## Funding

This research is based upon work supported in part by the Office of the Director of National Intelligence (ODNI), Intelligence Advanced Research Projects Activity (IARPA), via [2021-20120400001]. The views and conclusions contained herein are those of the authors and should not be interpreted as necessarily representing the official policies, either expressed or implied, of ODNI, IARPA, or the U.S. Government. The U.S. Government is authorized to reproduce and distribute reprints for governmental purposes notwithstanding any copyright annotation therein.

## Authors’ contributions

X.G.: conceptualization, data collection and curation, image analysis, investigation, methodology, writing—review and editing;

M.J.H.: conceptualization, writing—review and editing

N.H.F: conceptualization, project administration, supervision, and writing—review and editing.

N.P.-W.: conceptualization, formal analysis, investigation, methodology, project administration, supervision, writing—original draft and writing—review and editing.

## Notes

### Competing Interest Statement

The authors have declared no competing interest.

## References

Baltiansky, L., Frankel, G. and Feinerman, O. (2023). Emergent regulation of ant foraging frequency through a computationally inexpensive forager movement rule. Elife 12.

Baltiansky, L., Sarafian-Tamam, E., Greenwald, E. and Feinerman, O. (2021). Dual-fluorescence imaging and automated trophallaxis detection for studying multi-nutrient regulation in superorganisms. Methods in Ecology and Evolution 12, 1441–1457.

Barbee, B. and Pinter-Wollman, N. (2022). Nutritional needs and mortality risk combine to shape foraging decisions in ants. Current Zoology, zoac089.

Bates, D., Mächler, M., Bolker, B. M. and S.C., W. (2015). Fitting Linear Mixed-Effects Models using lme4. Journal of Statistical Software 67, 1–48.

Buffin, A., Denis, D., Van Simaeys, G., Goldman, S. and Deneubourg, J. L. (2009). Feeding and Stocking Up: Radio-Labelled Food Reveals Exchange Patterns in Ants. Plos One 4.

Cassill, D. L. and Tschinkel, W. R. (1995). Allocation of liquid food to larvae via trophallaxis in colonies of the fire ant, *Solenopsis invicta*. Animal Behaviour 50, 801–813.

Collier, M., Albery, G. F., McDonald, G. C. and Bansal, S. (2022). Pathogen transmission modes determine contact network structure, altering other pathogen characteristics. Proceedings of the Royal Society B-Biological Sciences 289.

Crall, J. D., Gravish, N., Mountcastle, A. M. and Combes, S. A. (2015). BEEtag: A Low-Cost, Image-Based Tracking System for the Study of Animal Behavior and Locomotion. Plos One 10.

Csardi, G. and Nepusz, T. (2006). The igraph software package for complex network research. International Journal of Complex Systems 1695, https://igraph.org.

Csata, E., Gautrais, J., Bach, A., Blanchet, J., Ferrante, J., Fournier, F., Levesque, T., Simpson, S. J. and Dussutour, A. (2020). Ant Foragers Compensate for the Nutritional Deficiencies in the Colony. Current Biology 30, 135-+.

Dahbi, A., Hefetz, A., Cerda, X. and Lenoir, A. (1999). Trophallaxis mediates uniformity of colony odor in Cataglyphis iberica ants (Hymenoptera, Formicidae). Journal of Insect Behavior 12, 559–567.

Dussutour, A. and Simpson, S. J. (2008). Carbohydrate regulation in relation to colony growth in ants. Journal of Experimental Biology 211, 2224–2232.

Dussutour, A. and Simpson, S. J. (2009). Communal Nutrition in Ants. Current Biology 19, 740–744.

Dussutour, A. and Simpson, S. J. (2012). Ant workers die young and colonies collapse when fed a high-protein diet. Proceedings of the Royal Society B-Biological Sciences 279, 2402–2408.

Eames, K. T. D. and Keeling, M. J. (2002). Modeling dynamic and network heterogeneities in the spread of sexually transmitted diseases. Proceedings of the National Academy of Sciences of the United States of America 99, 13330–13335.

Edelstein, A. D., Tsuchida, M. A., Amodaj, N., Pinkard, H., Vale, R. D. and Stuurman, N. (2014). Advanced methods of microscope control using muManager software. J Biol Methods 1.

Fefferman, N. H. and Ng, K. L. (2007). How disease models in static networks can fail to approximate disease in dynamic networks. Physical Review E 76, 031919.

Firth, J. A., Hellewell, J., Klepac, P., Kissler, S., Kucharski, A. J., Spurgin, L. G. and Grp, C. C.-W. (2020). Using a real-world network to model localized COVID-19 control strategies. Nature Medicine 26, 1616-+.

Fox, J. and Weisberg, S. (2019). An R Companion to Applied Regression. Third edition. Sage, *Thousand Oaks CA*..

Gates, M. C. and Woolhouse, M. E. J. (2015). Controlling infectious disease through the targeted manipulation of contact network structure. Epidemics 12, 11–19.

Godfrey, S. S. (2013). Networks and the ecology of parasite transmission: A framework for wildlife parasitology. Int J Parasitol Parasites Wildl 2, 235–45.

Gordon, D. M. (1989). Dynamics of task switching in harvester ants. Animal Behaviour 38, 194–204.

Gordon, D. M. (1996). The organization of work in social insect colonies. Nature 380, 121–124.

Gordon, D. M. (2010). Ant encounters: Interaction networks and colony behavior: Princeton University Press

Greene, M. J. and Gordon, D. M. (2006). Interaction rate informs harvester ant task decisions. Behavioral Ecology 18, 451–455.

Greene, M. J., Pinter-Wollman, N. and Gordon, D. M. (2013). Interactions with combined chemical cues inform harvester ant foragers’ decisions to leave the nest in search of food. Plos One 8, e52219.

Greenwald, E. E., Baltiansky, L. and Feinerman, O. (2018). Individual crop loads provide local control for collective food intake in ant colonies. Elife 7.

Hendriksma, H. P., Toth, A. L. and Shafir, S. (2019). Individual and Colony Level Foraging Decisions of Bumble Bees and Honey Bees in Relation to Balancing of Nutrient Needs. Frontiers in Ecology and Evolution 7.

Howard, D. F. and Tschinkel, W. R. (1980). The Effect of Colony Size and Starvation on Food Flow in the Fire Ant, Solenopsis-Invicta (Hymenoptera, Formicidae). Behavioral Ecology and Sociobiology 7, 293–300.

Howard, D. F. and Tschinkel, W. R. (1981). The Flow of Food in Colonies of the Fire Ant, Solenopsis-Invicta - a Multifactorial Study. Physiological Entomology 6, 297–306.

Jones, J. H. and Ready, E. (2022). The Emergent Structure of Subsistence Risk-Management Networks. American Journal of Biological Anthropology 177, 92–92.

Jones, K. L., Thompson, R. C. A. and Godfrey, S. S. (2018). Social networks: a tool for assessing the impact of perturbations on wildlife behaviour and implications for pathogen transmission. Behaviour 155, 689–730.

Kay, A. (2004). The relative availabilities of complementary resources affect the feeding preferences of ant colonies. Behavioral Ecology 15, 63–70.

Kohl, K. D., Coogan, S. C. P. and Raubenheimer, D. (2015). Do wild carnivores forage for prey or for nutrients?: Evidence for nutrient-specific foraging in vertebrate predators. Bioessays 37, 701–709.

Lenth, R. V. (2022). emmeans: Estimated Marginal Means, aka Least-Squares Means.. R package version *1.7.2.*, https://CRAN.R-project.org/package=emmeans.

Lloyd-Smith, J. O., Schreiber, S. J., Kopp, P. E. and Getz, W. M. (2005). Superspreading and the effect of individual variation on disease emergence. Nature 438, 355–359.

Lüdecke, D., Ben-Shachar, M. S., Patil, I., Waggoner, P. and Makowski, D. (2021). performance: An R Package for Assessment, Comparison and Testing of Statistical Models. Journal of Open Source Software 6, 3139.

Markin, G. P. (1970). Seasonal life cycle of Argentine ant, *Iridomyrmex-humilis* (Hymenoptera - Formicidae), in Southern California. Annals of the Entomological Society of America 63, 1238–1242.

Mayntz, D., Raubenheimer, D., Salomon, M., Toft, S. and Simpson, S. J. (2005). Nutrient-specific foraging in invertebrate predators. Science 307, 111–113.

Miller, J. S. and Pinter-Wollman, N. (2023). Social interactions differ in their impact on foraging decisions. Animal Behaivour.

Miller, J. S., Wan, E., O’Fallon, S. and Pinter-Wollman, N. (2022). Modularity and connectivity of nest structure scale with colony size. Evolution 76, 101–113.

O’Donnell, S. and Bulova, S. J. (2007). Worker connectivity: a simulation model of variation in worker communication and its effects on task performance. Insectes Sociaux 54, 211–218.

Pacala, S. W., Gordon, D. M. and Godfray, H. C. J. (1996). Effects of social group size on information transfer and task allocation. Evolutionary Ecology 10, 127–165.

Pautasso, M. and Jeger, M. J. (2008). Epidemic threshold and network structure: The interplay of probability of transmission and of persistence in small-size directed networks. Ecological Complexity 5, 1–8.

Pinter-Wollman, N. (2015a). Nest architecture shapes the collective behavior of harvester ants. Biology Letters, 20150695.

Pinter-Wollman, N. (2015b). Persistent variation in spatial behavior affects the structure and function of interaction networks. Current Zoology 61, 98–106.

Pinter-Wollman, N., Bala, A., Merrell, A., Queirolo, J., Stumpe, M. C., Holmes, S. and Gordon, D. M. (2013). Harvester ants use interactions to regulate forager activation and availability. Animal Behaviour 86, 197–207.

Pinter-Wollman, N., Wollman, R., Guetz, A., Holmes, S. and Gordon, D. M. (2011). The effect of individual variation on the structure and function of interaction networks in harvester ants. Journal of the Royal Society Interface 8, 1562–1573.

Portha, S., Deneubourg, J. L. and Detrain, C. (2002). Self-organized asymmetries in ant foraging: a functional response to food type and colony needs. Behavioral Ecology 13, 776–781.

Quque, M., Bles, O., Benard, A., Heraud, A., Meunier, B., Criscuolo, F., Deneubourg, J. L. and Sueur, C. (2021). Hierarchical networks of food exchange in the black garden antLasius niger. Insect Science 28, 825–838.

R Core, T. (2014). R: A language and environment for statistical computing. R Foundation for Statistical Computing, Vienna, Austria. URL http://www.R-project.org/.

Sah, P., Leu, S. T., Cross, P. C., Hudson, P. J. and Bansal, S. (2017). Unraveling the disease consequences and mechanisms of modular structure in animal social networks. Proceedings of the National Academy of Sciences of the United States of America 114, 4165–4170.

Salathé, M. and Jones, J. H. (2010). Dynamics and Control of Diseases in Networks with Community Structure. Plos Computational Biology 6, e1000736–e1000736.

Sendova-Franks, A. B., Hayward, R. K., Wulf, B., Klimek, T., James, R., Planque, R., Britton, N. F. and Franks, N. R. (2010). Emergency networking: famine relief in ant colonies. Animal Behaviour 79, 473–485.

Silk, M. J. and Fefferman, N. H. (2021). The role of social structure and dynamics in the maintenance of endemic disease. Behavioral Ecology and Sociobiology 75.

Simpson, S. J. and Raubenheimer, D. (2012). The nature of nutrition : a unifying framework from animal adaptation to human obesity. Princeton: Princeton University Press.

Sorensen, A. A. and Vinson, S. B. (1981). Quantitative Food Distribution Studies within Laboratory Colonies of the Imported Fire Ant, Solenopsis-Invicta Buren. Insectes Sociaux 28, 129–160.

Stroeymeyt, N., Casillas-Perez, B. and Cremer, S. (2014). Organisational immunity in social insects. Current Opinion in Insect Science 5, 1–15.

Udiani, O. and Fefferman, N. H. (2020). How disease constrains the evolution of social systems. Proceedings of the Royal Society B-Biological Sciences 287.

Ulrich, Y., Burns, D., Libbrecht, R. and Kronauer, D. J. C. (2016). Ant larvae regulate worker foraging behavior and ovarian activity in a dose-dependent manner. Behavioral Ecology and Sociobiology 70, 1011–1018.

Wallis, D. I. (1964). The foraging behaviour of the ant, Formica fusca. Behaviour 23, 149–176.

Wilson, E. O. and Eisner, T. (1957). Quantitative studies of liquid food transmission in ants. 4, 157–166.

